# Association of polygenic risk for major psychiatric illness with subcortical volumes and white matter integrity in UK Biobank

**DOI:** 10.1101/080283

**Authors:** LM Reus, X Shen, J Gibson, E Wigmore, L Ligthart, MJ Adams, G Davies, SR Cox, SP Hagenaars, ME Bastin, IJ Deary, HC Whalley, AM McIntosh

## Abstract

Major depressive disorder (MDD), schizophrenia (SCZ) and bipolar disorder (BP) are common, disabling and heritable psychiatric diseases with a complex overlapping polygenic architecture. Individuals with these disorders, as well as their unaffected relatives, show widespread structural differences in corticostriatal and limbic networks. Structural variation in many of these brain regions is also heritable and polygenic but whether their genetic architecture overlaps with major psychiatric disorders is unknown. We sought to address this issue by examining the impact of polygenic risk of MDD, SCZ, and BP on subcortical brain volumes and white matter (WM) microstructure in a large single sample of neuroimaging data; the UK Biobank Imaging study. The first release of UK Biobank imaging data compromised participants with overlapping genetic data and subcortical volumes (N = 978) and WM measures (N = 816). Our, findings however, indicated no statistically significant associations between either subcortical volumes or WM microstructure, and polygenic risk for MDD, SCZ or BP. In the current study, we found little or no evidence for genetic overlap between major psychiatric disorders and structural brain measures. These findings suggest that subcortical brain volumes and WM microstructure may not be closely linked to the genetic mechanisms of major psychiatric disorders.

## Introduction

Major depressive disorder (MDD), schizophrenia (SCZ) and bipolar disorder (BP) are major psychiatric disorders affecting between 1% and 13% of the general population^1,2^. Twin and family studies find a strong genetic contribution to all three of these disorders^3,4^ and genome-wide association studies (GWAS) suggest that these effects are conferred by the cumulative effect of many loci, each of small effect^5–7^. Whilst progress in identifying the specific risk-conferring loci in SCZ has proved highly productive^5^, progress in BD and MDD has been slower^6,7^. Several strategies may accelerate the discovery of risk-loci, including the use of quantitative traits such as brain structure and connectivity.

Brain structure and connectivity measures have been shown to be heritable quantitative traits, with high heritabilities reported for both subcortical volumes (44-88%)^8^ and white matter (WM) integrity (53-90%)^9^. Furthermore, several genetic variants that influence brain structure have been identified^10,11^. Subcortical brain volume and WM microstructure abnormalities in corticostriatal and limbic networks have also been shown to distinguish individuals with SCZ, BD and MDD from controls in several studies, albeit with some inconsistencies^12–17^. Differences between patients and controls in global measures of WM microstructure, diffusion tensor imaging (DTI) biomarkers fractional anisotropy (FA) and mean diffusivity (MD), have been more consistently reported in these disorders^18–23^, and could suggest wide-spread WM integrity reductions.

Together, these volumetric and WM brain differences potentially describe a network of abnormality associated with major psychiatric illness. This network may have an important mechanistic role in the development of these disorders. This possibility is supported by several studies that show an association between brain structure and genetic liability for psychiatric disorders, though due to the cost of imaging these samples are typically an order of magnitude smaller in size than the GWAS that preceded them. Notably, Van Scheltinga et al., (2013)^24^ found higher polygenic risk for SCZ to be modestly (R^2^ = 0.048) associated with smaller total brain volume, while Caseras et al., (2015)^25^ reported a negative association between polygenic risk scores (PGRS) for SCZ and BP, and globus pallidus and amygdala grey matter (GM) volumes. Similar results have been found for the association between less optimal WM microstructure and greater genetic liability for SCZ^24,26,27^.

These associations have not been consistently reported; specifically, several studies have found no association between subcortical volumes and polygenic risk for MDD and SCZ^28–30^. However, these studies have used differing methodologies (*ie.,* multi-scanner study^30^) with relatively small sample sizes (N^28^/N^29^ = 438/122), which may contribute to inconsistencies in findings.

The current study sought to investigate the genetic relationship between major mental disorders and structural brain measures in a single large quality-controlled dataset using the same scanner and same analysis pipelines. We used newly acquired data from the first release of the UK Biobank Imaging study to examine whether structural imaging measures, including subcortical volumes (N=978) and WM microstructure measures (N=816), were associated with genetic risk of MDD, SCZ, and BP^32^ (http://www.ukbiobank.ac.uk). We hypothesized that higher polygenetic risk for these disorders would be associated with decreased volumetric and WM integrity measures in specifically corticostriatal and limbic networks.

## Methods

### Study population

The first release of UK Biobank imaging data consisted of 4,446 subjects (N_females_/N_males_ = 2,342/2,104; mean age ± s.d. = 55.52 ± 7.62 years) with quality-checked volumetric and DTI data for the analysis (conducted by UK Biobank, Brain Imaging Documentation V1.1, http://www.ukbiobank.ac.uk)^33^. In the current study, subjects were excluded if they did not provide genetic data, or if they were known to have participated in studies from the Psychiatric Genomics Consortium (PGC) GWAS dataset as this was used to construct the polygenic risk scores (PGRS). In total, the volumetric analysis included 978 subjects (N_females_/N_males_ = 499/479; mean age ± s.d. = 55.32 ± 7.38 years) and the DTI analysis included 816 subjects (N_females_/N_males_ = 413/403; mean age ± s.d. = 55.49 ± 7.26 years). In addition, a sensitivity analysis was performed where we imposed internal imaging quality checks excluding individuals whose subcortical volume or FA values were greater than three standard deviations above or below the mean for that measure. The sensitivity analysis, further referred as the sample excluding outliers, included 892 (N_females_/N_males_ = 477/415; mean age ± s.d. = 55.29 ± 7.31 years) and 733 (N_females_/N_males_ = 373/360; mean age ± s.d. = 55.31 ± 7.22 years) subjects for the volumetric and DTI analysis respectively.

This study has been approved by the National Health Service (NHS) Research Ethics Service (approval letter dated 17^th^ June 2011, reference: 11/NW/0382), and by the UK Biobank Access Committee (Project #4844). Written informed consent was obtained from each subject.

### Genotyping and derivation of polygenic risk scores

Procedures for DNA collection and genotyping in UK Biobank have been described previously by Hagenaars et al., (2016)^34^. Subjects were excluded from the PGRS analysis if they had a non-British ancestry, gender mismatch, relatedness (r>0.044), and genotype missingness bigger than 2%.

PGRS for the three traits of interest (*i.e.,* MDD, SCZ, BP) were created using PRSice^35^. For each subject PGRS were calculated by adding the sum of each allele weighted by the strength of its association with the trait of interest. The strength of this association has been calculated previously by the PGC^5–7,36^. Prior to calculating the PGRS, single nucleotide polymorphisms (SNPs) were excluded if they had a minor allele frequency less than 1%, deviated significantly from Hardy-Weinberg equilibrium (p < 1x10^-6^) in the total sample of founder individuals, or had a call rate of less than 99%. The remaining SNPs were used to calculate 15 multidimensional scaling (MDS) ancestry components (to account for population structure). For the main results, the SNP inclusion threshold was set as p ≤ 0.5, as Purcell et al., (2009)^37^ and studies following this work, have shown that this threshold is generally the most predictive of case-control status in subsequent studies^5–7,36^. Using the p ≤ 0.5 SNP inclusion threshold, 631,763, 5,422,836 and 1,283,324 SNPs (after LD pruning) were included for the calculation of MDD-PGRS, SCZ-PGRS and BP-PGRS, respectively. Results generated by a SNP inclusion threshold of p ≤ 0.01, p ≤ 0.05, p ≤ 0.1 and p ≤ 1.0 are reported in supplementary materials.

### Image acquisition and Neuroimaging data pre-processing

Procedures for image acquisition and pre-processing are available on the UK Biobank website (http://www.ukbiobank.ac.uk/), and have been documented previously^38^. More detailed information is reported in the supplementary materials.

### Statistical analysis

All analyses were performed using R (version 3.2.3) in the Linux environment (R Development Core Team, 2010). To examine whether higher PGRS were associated with globally poor WM microstructure, general components for FA (*g*FA) and MD (*g*MD) across the 27 WM tracts were calculated^38^. Latent measures, explaining a portion of variance WM structure, were calculated for both FA and MD by extracting the scores on the first unrotated components, using a principal component analysis (PCA) on the 27 WM tracts. The first components accounted for a mean of 44.82% and 41.73% of total variance in FA and, 47.45% and 45.35% in MD, in the sample including and excluding outliers respectively (loadings of tracts on the first latent component > 0.15). To examine whether higher PGRS were selectively associated with poor WM structure in a certain classification, this PCA was also performed separately for groups of different categories of WM tracts, including association (inferior fronto-occipital fasciculus, inferior longitudinal fasciculus, superior longitudinal fasciculus, uncinate fasciculus, cingulum), projection (forceps minor, forceps major, corticospinal tracts, acoustic radiation, medial lemniscus, middle cerebellar peduncle) and thalamic radiation fibers, separately^38^ (sample including outliers: variance explained = 54.52%, 38.68% and 61.92% respectively for each subset, and loading of first component > 0.33; sample excluding outliers: variance explained = 60.05%, 32.05% and 72.05% respectively for each subset and loading of first component > 0.27).

For bilateral brain regions, associations between the three PGRSs and each imaging measure were examined using a repeated measure linear mixed-effects analysis, modelling hemisphere as a random factor (where the PGRS * hemisphere interaction itself was found to be non-significant). For unilateral structures (*e.g.* total GM volume, forceps major, forceps minor and middle cerebellar peduncle), and those where there was a significant interaction between PGRS and hemisphere, associations were analysed as single structures without repeated measurements and without hemisphere as a separate term in the model.

For all models, additional fixed effects included age, age^2^, gender, genotype batch, genotype array and 15 MDS components. Intracranial volume, calculated as the sum of WM, GM and ventricular CSF, was also modelled as a fixed effect in the analysis of subcortical volumes in order to adjust for differences in overall brain size. PGRS and imaging measures were scaled to zero mean and unitary standard deviation. Standardised Beta values are reported throughout results. Results were corrected for multiple comparisons using a false discovery rate (FDR) correction^39^. Nagelkerke's R^2^ was estimated to assess the proportion of variation explained by PGRS.

Post-hoc positive control analyses, with age as an independent variable and each of the brain measures as dependent variable, were performed in order to test the sensitivity of UK Biobank imaging, by verifying that we could replicate the widely reported negative associations between many structural brain measures and age (including those reported previously in an overlapping sample in UK Biobank^38^). Fixed effects included gender for all structures, and side of hemisphere for lateralized structures only.

## Results

### PGRS and total brain and subcortical volumes

No significant associations were observed between PGRS (MDD, SCZ, or BP) at the threshold of p ≤ 0.5 and total GM, WM or CSF volume, in samples including or excluding outliers (Table 1). Furthermore, no significant associations were found between any of the three sets of PGRS and total GM, WM and CSF volume for any of the other PGRS p-value thresholds (all β < 0.054; all p_uncorrected_ and p FDR > 0.05) (Table S1).

**Table 1.** Results - PGRS and total brain and subcortical volumes. Association of PGRS (MDD, SCZ or BP) at p ≤ 0.5 with total grey matter, white matter, cerebrospinal fluid, and subcortical volumes, in sample including and excluding outliers.

|  | Including outliers (N = 978) |  |  |  |  | Excluding outliers (N = 892) |  |  |  |  |
| --- | --- | --- | --- | --- | --- | --- | --- | --- | --- | --- |
|  | Beta: z ratio (S.D.) | t statistic | p -uncorr. | p-FDR | R<sup>2</sup> | Beta: z ratio (S.D.) | t statistic | p -uncorr | p-FDR | R<sup>2</sup> |
| **MDD-PGRS** |  |  |  |  |  |  |  |  |  |  |
| GM volume | 0.001 (0.027) | 0.049 | 0.961 | 1.000 | 0.000 | -0.001 (0.029) | -0.051 | 0.959 | 1.000 | 0.000 |
| WM volume | -0.007 (0.027) | -0.262 | 0.793 | 1.000 | 0.005 | -0.026 (0.028) | -0.913 | 0.362 | 1.000 | 0.067 |
| CSF volume | 0.042 (0.030) | 1.392 | 0.164 | 0.492 | 0.174 | 0.054 (0.031) | 1.748 | 0.081 | 0.243 | 0.289 |
| Caudate | 0.026 (0.026) | 0.970 | 0.332 | 1.000 | 0.066 | 0.031 (0.028) | 1.119 | 0.264 | 1.000 | 0.099 |
| Hippocampus | -0.002 (0.026) | -0.082 | 0.935 | 1.000 | 0.000 | -0.003 (0.026) | -0.130 | 0.896 | 1.000 | 0.001 |
| Pallidum | 0.003 (0.027) | 0.107 | 0.915 | 1.000 | 0.001 | -0.013 (0.027) | -0.466 | 0.641 | 1.000 | 0.016 |
| Thalamus | -0.012 (0.020) | -0.611 | 0.541 | 1.000 | 0.015 | -0.025 (0.020) | -1.218 | 0.223 | 1.000 | 0.060 |
| Amygdala | -0.000 (0.027) | -0.010 | 0.992 | 1.000 | 0.000 | 0.015 (0.027) | 0.551 | 0.582 | 1.000 | 0.023 |
| Nucleus accumbens | -0.019 (0.026) | -0.742 | 0.458 | 1.000 | 0.038 | -0.030 (0.027) | -1.126 | 0.260 | 1.000 | 0.093 |
| Putamen | 0.007 (0.022) | 0.309 | 0.757 | 1.000 | 0.005 | -0.010 (0.023) | -0.451 | 0.652 | 1.000 | 0.011 |
| **SCZ-PGRS** |  |  |  |  |  |  |  |  |  |  |
| GM volume | 0.003 (0.027) | 0.093 | 0.926 | 1.000 | 0.001 | -0.018 (0.029) | -0.613 | 0.540 | 1.000 | 0.033 |
| WM volume | 0.000 (0.026) | 0.019 | 0.985 | 1.000 | 0.000 | -0.004 (0.029) | -0.147 | 0.883 | 1.000 | 0.002 |
| CSF volume | -0.007 (0.029) | -0.246 | 0.805 | 1.000 | 0.005 | 0.063 (0.031) | 2.029 | 0.043 | 0.128 | 0.397 |
| Caudate | -0.003 (0.026) | -0.101 | 0.919 | 1.000 | 0.001 | -0.016 (0.028) | -0.559 | 0.577 | 1.000 | 0.025 |
| Hippocampus | -0.001 (0.025) | -0.044 | 0.965 | 1.000 | 0.000 | 0.015 (0.026) | 0.558 | 0.577 | 1.000 | 0.021 |
| Pallidum | -0.015 (0.027) | -0.578 | 0.564 | 1.000 | 0.024 | -0.022 (0.027) | -0.795 | 0.427 | 1.000 | 0.047 |
| Thalamus | -0.039 (0.020) | -1.929 | 0.054 | 0.378 | 0.149 | -0.039 (0.020) | -1.947 | 0.052 | 0.363 | 0.155 |
| Amygdala | 0.015 (0.026) | 0.558 | 0.577 | 1.000 | 0.022 | 0.025 (0.028) | 0.912 | 0.362 | 1.000 | 0.063 |
| Nucleus accumbens | -0.024 (0.026) | -0.924 | 0.356 | 1.000 | 0.058 | -0.023 (0.027) | -0.843 | 0.400 | 1.000 | 0.053 |
| Putamen | 0.005 (0.022) | 0.211 | 0.833 | 1.000 | 0.002 | -0.016 (0.023) | -0.672 | 0.502 | 1.000 | 0.025 |

| BP-PGRS |  |  |  |  |  |  |  |  |  |  |
| --- | --- | --- | --- | --- | --- | --- | --- | --- | --- | --- |
| **GM volume** | 0.003 (0.027) | 0.093 | 0.926 | 1.000 | 0.001 | -0.012 (0.029) | -0.405 | 0.686 | 1.000 | 0.013 |
| **WM volume** | 0.000 (0.026) | 0.019 | 0.985 | 1.000 | 0.000 | 0.006 (0.028) | 0.229 | 0.819 | 1.000 | 0.004 |
| **CSF volume** | -0.007 (0.029) | -0.246 | 0.805 | 1.000 | 0.005 | 0.007 (0.030) | 0.216 | 0.829 | 1.000 | 0.004 |
| **Caudate** | 0.017 (0.026) | 0.653 | 0.514 | 1.000 | 0.029 | 0.007 (0.028) | 0.245 | 0.806 | 1.000 | 0.005 |
| **Hippocampus** | 0.024 (0.025) | 0.963 | 0.336 | 1.000 | 0.058 | 0.011 (0.026) | 0.421 | 0.674 | 1.000 | 0.012 |
| **Pallidum** | -0.000 (0.026) | -0.009 | 0.993 | 1.000 | 0.000 | -0.023 (0.027) | -0.865 | 0.387 | 1.000 | 0.054 |
| **Thalamus** | 0.010 (0.020) | 0.517 | 0.605 | 1.000 | 0.010 | 0.005 (0.020) | 0.249 | 0.803 | 1.000 | 0.002 |
| **Amygdala** | 0.030 (0.026) | 1.158 | 0.247 | 1.000 | 0.091 | 0.032 (0.027) | 1.186 | 0.236 | 1.000 | 0.103 |
| **Nucleus accumbens** | -0.008 (0.026) | -0.296 | 0.767 | 1.000 | 0.006 | -0.017 (0.027) | -0.626 | 0.532 | 1.000 | 0.028 |
| **Putamen** | 0.025 (0.022) | 1.133 | 0.257 | 1.000 | 0.061 | 0.015 (0.023) | 0.659 | 0.510 | 1.000 | 0.023 |

MDD: major depressive disorder, SCZ: schizophrenia, BP: bipolar disorder, PGRS: polygenic risk scores, uncorr.: uncorrected, FDR: false discovery rate, GM: grey matter, WM: white matter, CSF: cerebrospinal fluid, S.D.: standard deviation. Controlled for age, age<sup>2</sup>, gender, genotype batch and array, and 15 MDS components. Additional fixed effects for subcortical volumes include intracranial volume and side of hemisphere. R<sup>2</sup> = estimate of variance explained by PGRS in %.

A modest negative association was observed between thalamus volumes and the polygenic risk for SCZ at the threshold of p ≤ 0.05, p ≤ 0.1, p ≤ 0.5 and p ≤ 1.0 (β range = −0.043 – −0.039; p_uncorrected_ = 0.052, p_uncorrected_ = 0.030, p_uncorrected_ = 0.054, p_uncorrected_ = 0.048 for all PGRS thresholds respectively) (Table 1 and S3). However, following FDR-correction no significant associations were found between PGRS (MDD, SCZ, or BP) at the threshold of p ≤ 0.5 and all subcortical volumes, before and after exclusion of outliers (Table 1). These results were unchanged at the other p-value thresholds (Table S2, S3, S4). For subcortical volumetric measures, no significant PGRS * hemisphere interactions were found; therefore, all analyses of bilateral structures were conducted using a repeated measures design with hemisphere as a repeated factor.

### PGRS and diffusion measures

Associations between PGRS (MDD, SCZ, BP) and measures of general WM microstructure (*g*FA and *g*MD) were not significant (*g*FA β range = −0.048 – 0.017; p range = 0.216 – 0.771, *g*MD β range = −0.047 – 0.031; p range = 0.181 – 0.985) (Table 2 and 3). Moreover, when *g*FA and *g*MD were calculated separately for projection, association, and thalamic radiation fibres, there remained no signification association between these measures and PGRS for MDD, SCZ and BP at p ≤ 0.5 threshold (Table 2 and 3). These findings were similar at other p-value thresholds (Table S5 and S6).

**Table 2.** Results - PGRS and diffusion measures. Association of PGRS (MDD, SCZ or BP) at p ≤ 0.5 with *g*FA and *g*FA for association, projection and thalamic WM fibers.

|  | Including outliers (N = 816) |  |  |  | Excluding outliers (N = 733) |  |  |  |
| --- | --- | --- | --- | --- | --- | --- | --- | --- |
|  | Beta: z ratio (S.D.) | t statistic | p | R<sup>2</sup> | Beta: z ratio (S.D.) | t statistic | p | R<sup>2</sup> |
| **MDD-PGRS** |  |  |  |  |  |  |  |  |
| *gFA* | 0.017 (0.036) | 0.476 | 0.634 | 0.030 | -0.011 (0.038) | -0.291 | 0.771 | 0.012 |
| *gFA* Association | 0.037 (0.036) | 1.026 | 0.305 | 0.139 | 0.011 (0.039) | 0.296 | 0.767 | 0.013 |
| *gFA* Projection | -0.020 (0.036) | -0.560 | 0.576 | 0.041 | -0.049 (0.038) | -1.297 | 0.195 | 0.244 |
| *gFA* Thalamic | 0.002 (0.037) | -0.043 | 0.966 | 0.000 | -0.025 (0.039) | -0.650 | 0.516 | 0.063 |
| **SCZ-PGRS** |  |  |  |  |  |  |  |  |
| *gFA* | -0.028 (0.036) | -0.768 | 0.443 | 0.078 | -0.048 (0.039) | -1.237 | 0.216 | 0.230 |
| *gFA* Association | -0.020 (0.036) | -0.559 | 0.576 | 0.041 | -0.040 (0.039) | -1.034 | 0.302 | 0.161 |
| *gFA* Projection | -0.034 (0.036) | -0.947 | 0.344 | 0.116 | -0.036 (0.038) | -0.947 | 0.344 | 0.132 |
| *gFA* Thalamic | -0.032 (0.037) | 0.885 | 0.376 | 0.105 | -0.059 (0.039) | -1.521 | 0.129 | 0.348 |
| **BP-PGRS** |  |  |  |  |  |  |  |  |
| *gFA* | -0.013 (0.036) | -0.367 | 0.714 | 0.017 | -0.014 (0.038) | -0.370 | 0.711 | 0.019 |
| *gFA* Association | -0.016 (0.036) | -0.442 | 0.659 | 0.025 | -0.017 (0.038) | -0.445 | 0.657 | 0.028 |
| *gFA* Projection | -0.023 (0.035) | -0.665 | 0.506 | 0.055 | -0.014 (0.037) | -0.369 | 0.712 | 0.019 |
| *gFA* Thalamic | -0.000 (0.036) | 0.0223 | 0.982 | 0.000 | -0.008 (0.038) | -0.214 | 0.831 | 0.008 |

FA: fractional anisotropy, WM: white matter, g: general component, MDD: major depressive disorder, SCZ: schizophrenia, BP: bipolar disorder, PGRS: polygenic risk scores, S.D.: standard deviation. Controlled for age, age<sup>2</sup> and gender, genotype batch and array, and 15 MDS components. R<sup>2</sup> = estimate of variance explained by PGRS in %.

**Table 3.** Association of PGRS (MDD, SCZ or BP) at p ≤ 0.5 with *g*MD and *g*MD for association, projection and thalamic WM fibers.

|  | Including outliers (N = 816) |  |  |  | Excluding outliers (N = 733) |  |  |  |
| --- | --- | --- | --- | --- | --- | --- | --- | --- |
|  | Beta: z ratio (S.D.) | t statistic | p | R<sup>2</sup> | Beta: z ratio (S.D.) | t statistic | p | R<sup>2</sup> |
| **MDD-PGRS** |  |  |  |  |  |  |  |  |
| *gMD* | -0.047 (0.035) | -1.340 | 0.181 | 0.224 | -0.028 (0.037) | -0.756 | 0.450 | 0.079 |
| *gMD* Association | -0.053 (0.036) | -1.484 | 0.138 | 0.281 | -0.030 (0.038) | -0.810 | 0.418 | 0.093 |
| *gMD* Projection | -0.053 (0.037) | -1.417 | 0.157 | 0.277 | -0.042 (0.039) | -1.075 | 0.283 | 0.179 |
| *gMD* Thalamic | -0.031 (0.034) | -0.907 | 0.365 | 0.094 | -0.017 (0.035) | -0.479 | 0.632 | 0.029 |
| **SCZ-PGRS** |  |  |  |  |  |  |  |  |
| *gMD* | -0.018 (0.035) | -0.503 | 0.615 | 0.031 | 0.001 (0.037) | 0.019 | 0.985 | 0.000 |
| *gMD* Association | -0.006 (0.036) | -0.179 | 0.858 | 0.004 | 0.005 (0.038) | 0.121 | 0.903 | 0.002 |
| *gMD* Projection | -0.035 (0.037) | -0.951 | 0.342 | 0.124 | -0.021 (0.040) | -0.517 | 0.606 | 0.042 |
| *gMD* Thalamic | -0.026 (0.034) | -0.777 | 0.438 | 0.068 | 0.003 (0.036) | 0.084 | 0.933 | 0.001 |
| **BP-PGRS** |  |  |  |  |  |  |  |  |
| *gMD* | 0.014 (0.034) | 0.408 | 0.684 | 0.020 | 0.031 (0.036) | 0.865 | 0.387 | 0.099 |
| *gMD* Association | 0.017 (0.035) | 0.500 | 0.617 | 0.030 | 0.032 (0.037) | 0.874 | 0.383 | 0.104 |
| *gMD* Projection | 0.003 (0.036) | 0.082 | 0.935 | 0.000 | -0.009 (0.039) | 0.232 | 0.817 | 0.008 |
| *gFMD*Thalamic | 0.011 (0.033) | 0.328 | 0.743 | 0.012 | 0.032 (0.035) | 0.933 | 0.351 | 0.104 |

MD: mean diffusivity, WM: white matter, g: general factor, MDD: major depressive disorder, SCZ: schizophrenia, BP: bipolar disorder, PGRS: polygenic risk scores, S.D.: standard deviation. Controlled for age, age<sup>2</sup> and gender, genotype batch and array, and 15 MDS components. R<sup>2</sup> = estimate of variance explained by PGRS in %.

Trendwise associations were observed between some polygenic MDD risk scores and MD values in the anterior thalamic radiation, a group of WM fibers connecting frontal regions to anterior and middle nuclear groups of the thalamus (β = 0.084; p-FDR = 0.085, β = 0.088; p-FDR = 0.061, β = 0.086; p-FDR = 0.065 for PGRS threshold p ≤ 0.01, p ≤

0.05 and p ≤ 0.5 respectively) (Table S9, S10). Polygenic MDD, SCZ or BP risk was also not significantly associated with other individual tract-specific water diffusion measures in the sample including (N = 816) and excluding outliers (N = 733) (Table S7 and S9). These results were again similar for PGRS at the other thresholds (Table S8 and S10).

No significant PGRS * hemisphere interactions were found for the majority of individual tract water diffusion values, and therefore analyses on bilateral structures were conducted as a repeated measure. However, the SCZ-PGRS * hemisphere interaction was found to be significant for FA values in the parahippocampal portion of the cingulum (connecting orbitofrontal regions to the hippocampus) in the sample including outliers (β = −0.052; p_corrected_ = 0.032). We therefore conducted tests of this association with SCZ-PGRS separately for each lateralised structure for this region. However, these were also not significant for FA in either the left (β = −0.040; p_uncorrected_ = 0.262) or right (β = −0.079; p_uncorrected_ = 0.029) parahippocampal portion of the cingulum, following FDR correction.

### Age-related effects in structural brain measures

We additionally performed a post-hoc validation analysis in order to verify that despite our lack of polygenic associations we could indeed replicate the widely reported negative associations between structural brain measures and age. We conducted our analyses using age as an independent variable and each of the brain measures as dependent variable. As expected, total GM and WM volumes showed a significant negative association with age, whereas this association was positive for total CSF volume (Table S11). Additionally, highly significant (β = −0.392 - −0.166; p-FDR < 0.001) negative associations were reported between age and the majority of subcortical volumes. Amygdala volumes were not significant associated with age (Table S12).

The association between age and WM microstructure has been examined in a larger sample (N = 3,513) of the UK Biobank dataset^38^. Age effects on WM in this sample with overlapping MRI and genetic data are broadly in agreement with this previous study, as indicated by a negative association between age and WM microstructure quality (Table S13 and S14).

## Discussion

Structural brain abnormalities in corticostriatal and limbic structures are thought to play an important role in the pathophysiology of common disabling psychiatric disorders, such as MDD, SCZ and BP. Since these structural brain abnormalities have been observed in unaffected relatives of patients and have shown to be heritable quantitative traits, the current study sought to test the impact of the polygenic liability for MDD, SCZ, or BP on subcortical brain volumes and WM microstructure in individuals from a population-based study. Contrary to our predictions, we found no evidence for an association between polygenic risk for MDD, SCZ and BP and either subcortical brain regions or WM microstructure. These findings provide no support for the hypothesis that the polygenic liability for MDD, SCZ and BP is linked to structural brain changes in major psychiatric illness.

Our findings are in contrast to many published studies that report a relationship between subcortical GM volume and PGRS for psychiatric disorders^24,25^. However, these studies have generally been smaller than UK Biobank (N^24^/N^35^ = 294/274), increasing the risk of false positive results. In a complementary approach to our study, schizophrenia polygenic risk scores and linkage disequilibrium (LD) score regression have been used to test for shared genetic architecture between subcortical brain volumes and schizophrenia (N = 11,840)^30^. As in the current study, this study also reported a lack of overlapping genetic architecture between SCZ and subcortical volumes^30^. Moreover, Van Scheltinga et al.’s^24^ result of a significant association between and polygenic liability for SCZ and total brain volume could not be replicated in another study^29^. Holmes et al., (2012) (N = 438) also reported no association between MDD-PGRS and amygdala volume^28^. Here we extend these findings to SCZ, BD and MDD and across a range of subcortical measures (Table 1, 2, and S1-S4) using a large single scanner sample.

Only a few studies have been published regarding the association between genetic liability for psychiatric illness and WM integrity. No associations between WM integrity and polygenetic liability have been reported for BP, whereas polygenetic liability for MDD and SCZ was associated with decreased WM integrity^26,27^. Furthermore, individuals at high genetic risk for psychiatric disorders, such as relatives of patients, have shown reduced WM integrity^40–42^. Conversely, our study found no relation between genetic liability for psychiatric disorders and WM integrity (Table 3-5, and S5-S10).

The present study has some important limitations that should be taken into account. Firstly, it is possible that any shared genetic architecture with brain structure was too small to detect in the current sample. Although our sample sizes (N_volumetric/_N_DTI_ = 978/816) are large for imaging research, they are relatively small when compared to a typical PGRS study. Moreover, our sample is a mixed community population, which included both healthy individuals, patients, and individuals without clinical records. To exclude the possibility of different genetic associations across clinical groups, future research should replicate our findings in cases and controls, separately. A general limitation of PGRS is that they account for only a small proportion of total phenotypic variance, far short of their narrow sense heritability. The addition of further discovery GWA samples for MDD, SCZ and BD will improve the predictive accuracy of PGRS and alongside developments in imaging consortia, such as ENIGMA (http://enigma.ini.usc.edu/), will make it possible to test for smaller effects in future studies^43^. The current results indicate that polygenic risk score associations with brain structure should be interpreted cautiously in smaller studies. Our findings should, however, be replicated to exclude the presence of a small genetic correlation between psychiatric disorder and brain structure and connectivity measures. Furthermore, future work should also examine whether the genetic architecture of other *in vivo* brain parameters, such as measures of cortical thickness and brain function overlap with major mental illness.

In summary, the current study reports no evidence of shared genetic architecture between psychiatric disorders and either subcortical brain volumes or WM integrity. Our results do not replicate the findings of several published studies with small sample sizes, but are supported by similar recent work, particularly in SCZ.

## Acknowledgements

This work was supported by a grant of the Wellcome Trust Strategic Award “Stratifying Resilience and Depression Longitudinally’’ (STRADL) (reference 104036/Z/14/Z).

We thank the UK Biobank participants for their participation, and the UK Biobank team for their work in collecting and providing these data for analysis. Part of the work was undertaken in The University of Edinburgh Centre for Cognitive Ageing and Cognitive Epidemiology (CCACE), part of the cross council Lifelong Health and Wellbeing Initiative (MR/K026992/1); funding from the Biotechnology and Biological Sciences Research Council (BBSRC) and Medical Research Council (MRC) is gratefully acknowledged. Age UK (The Disconnected Mind project) also provided support for the work undertaken at CCACE.

LMR was supported by an Erasmus Traineeship Grant. XS receives support from China Scholarship Council. HCW is supported by a JMAS SIM fellowship from the Royal College of Physicians of Edinburgh and by an ESAT College Fellowship from the University of Edinburgh. SRC was supported by MRC grant MR/M013111/1. IJD is supported by the Medical Research Council award to CCACE (MR/K026992/1) and by the Dementias Platform UK (MR/L015382/1). IJD and SRC are additionally supported by the Age UK-funded Disconnected Mind project (http://www.disconnectedmind.ed.ac.uk).

## Author Contributions

AMM, HCW, SRC, XS and LMR contributed to the design of the study, analysis of the data, and writing the manuscript. MJA was involved in curating the data. IJD and MEB were involved in the conception of the study and overseeing analysis methodology. JG, EW and SPH were involved in the preprocessing and calculation of the polygenic risk scores. UK Biobank collected all data and was involved in the preprocessing of imaging data. All author discussed and commented on the manuscript.

## Competing interests

AMM has previously received grant support from Pfizer, Lilly and Janssen. These studies are not connected to the current investigation. Remaining authors report no competing financial interests.

## Corresponding author

Correspondence to Dr. Heather C. Whalley.

